# Single-cell analysis of the nervous system at small and large scales with instant partitions

**DOI:** 10.1101/2023.07.14.549051

**Authors:** PW Frazel, K Fricano-Kugler, AA May-Zhang, MR O’Dea, P Prakash, NM Desmet, H Lee, RH Meltzer, KM Fontanez, P Hettige, Y Agam, G Lithwick-Yanai, D Lipson, BW Luikart, JD Dasen, SA Liddelow

## Abstract

Single-cell RNA sequencing is a new frontier across all biology, particularly in neuroscience. While powerful for answering numerous neuroscience questions, limitations in sample input size, and initial capital outlay can exclude some researchers from its application. Here, we tested a recently introduced method for scRNAseq across diverse scales and neuroscience experiments. We benchmarked against a major current scRNAseq technology and found that PIPseq performed similarly, in line with earlier benchmarking data. Across dozens of samples, PIPseq recovered many brain cell types at small and large scales (1,000-100,000 cells/sample) and was able to detect differentially expressed genes in an inflammation paradigm. Similarly, PIPseq could detect expected and new differentially expressed genes in a brain single cell suspension from a knockout mouse model; it could also detect rare, virally-labelled cells following lentiviral targeting and gene knockdown. Finally, we used PIPseq to investigate gene expression in a nontraditional model species, the little skate (Leucoraja erinacea). In total, PIPSeq was able to detect single-cell gene expression changes across models and species, with an added benefit of large scale capture and sequencing of each sample.

## INTRODUCTION

The brain is a richly heterogenous tissue, that requires a symphony of interactions between cell types to maintain homeostasis and for proper functioning. Single cell RNA sequencing (scRNAseq) has unlocked cell type heterogeneity during normal physiology^1^, including development and aging, in addition to during many neurological diseases. These single cell sequencing datasets have allowed for unprecedented, million-cell resolution of diversity across the mature mammalian brain^2,3^. These atlases have begun to uncover fundamental cell type identities across the brain^4^, including subtle but important differences across transcriptionally similar cell types such as astrocytes^5^. Similarly, these datasets have uncovered key subtypes of cells driving disease states in neurodegeneration and neurodevelopment^6,7^. The next era of scRNAseq studies will allow researchers to profile transient states that inform the mature cell type^7^ or disease state of interest.

While initial atlases of individual cell/cell population gene expression have uncovered many new hypotheses, these experiments will always face the need to enrich the subtype of interest (impossible for exploratory experiments) or sidestep this by sampling at large scale. The need to capture rare cells and subtypes has led to the proliferation of cell atlases of the developing^8-10^ and adult brain^11,12^. Single cell methods diverge greatly in terms of technology and throughput^13^, and are a topic of great current interest across neuroscience^14^. Sample prep can be hard for brain tissue, particularly heavily myelinated regions (like the spinal cord and corpus callosum) and peripheral tissue types that contain excess connective tissue has proven difficult to isolate high quality cells for capture and sequencing^15^. A second, equally pressing problem has been the difficulty in capture of some cell types – for example, astrocytes – which leads to the gross underrepresentation of these cells across most datasets^16^. Aged brain has also been difficult for isolation and capture of single cells with buildup of fatty tissues and myelin that making sample preparation for scRNAseq difficult and unreliable^15,17^. These excesses of non-cellular components and debris often require multiple rounds of ‘clean up’ using FACS or magnetic beads so that microfluidic-based collection methods do not clog. This additional time required for sample preparation can then lead to gene expression artifacts^15^.

Here we use a microfluidic-free partition^18^ method for single cell RNA sequencing (scRNAseq) of nervous system samples using PIPseq (Particle-Templated Instant Partitions sequencing) that could potentially help with both problems (cell capture and scalability). We tested PIPseq across a broad range of nervous tissue and experimental types to assess its utility, with comparable results to microfluidics-based scRNAseq (10X Genomics) as has been reported already^18^.

The datasets here span two species, sample age (young to old), input tissue preparation (fresh, frozen, shipped), cells and nuclei, as well as compatibility with multiple sequencing platforms (Illumina vs. Ultima). Overall, these data demonstrate that PIPseq is a low-cost, scalable, and easily accessible method for any researchers seeking to complete sc/ snRNAseq studies.

## RESULTS

### Direct comparison of PIPseq scRNAseqto 10X Genomics

We first wanted to ensure that the quality and depth of sequencing provided by PIPseq was comparable to other commercial microfluidic-based scRNAseq technologies, namely the 10X Genomics Next GEM v3.1 platform. We analyzed dissociated cells from the same samples collected from young (postnatal day, P30) mouse brains using both PIPseq and 10X Genomics platforms to directly compare the two methods. Mouse forebrain was dissociated into single-cell suspension using a standard papain-based approach^19^, and 40,000 cells were loaded into v2.1 PIPs and captured after 135 seconds of vortexing (following manufacturer’s instructions). In parallel, ∼34,000 cells per sample from this same prep were loaded onto a 10X Genomics microfluidics chip for analysis with the 3’ Gene Expression kit (Fig 1A, Fig S1). As expected, each technique yielded different numbers of recovered cells at different sequencing depths (21,203 cells for PIPseq and 24,562 cells for 10X) so we randomly downsampled the FASTQs for each sample to the same read depth (237.5 million reads) and used the top 21,203 cells from each dataset (11,200 reads/cell), to compare equal numbers of cells across the two platforms and minimize the inclusion of non-cell barcodes that were called using the default CellRanger v6 algorithm. Once equally downsampled, we noted broad mapping and quality-control concordance between PIPseq and 10x (Fig S1A,B), with some interesting differences. We detected a higher proportion of PIPseq reads mapped to the mouse transcriptome compared to 10X (72% mapping [PIP-seq] vs. 50.4% mapping [10X]). Conversely, we observed higher median genes per cell in 10X data (778 [PIP-seq] vs. 1254 [10X]), which was mirrored by increased median UMIs per cell in 10X vs. PIPseq (1106 [PIP-seq] vs. 2081 [10X]). Relatedly, sequencing saturation was higher in PIPseq vs. 10X at the same downsampled read depth (0.37 [PIP-seq] vs. 0.17 [10X], suggesting lower complexity in the PIPseq library. We note that the two platforms use different read mapping algorithms (CellRanger vs. PIPSeeker), which complicates these comparisons.

**Figure 1.**
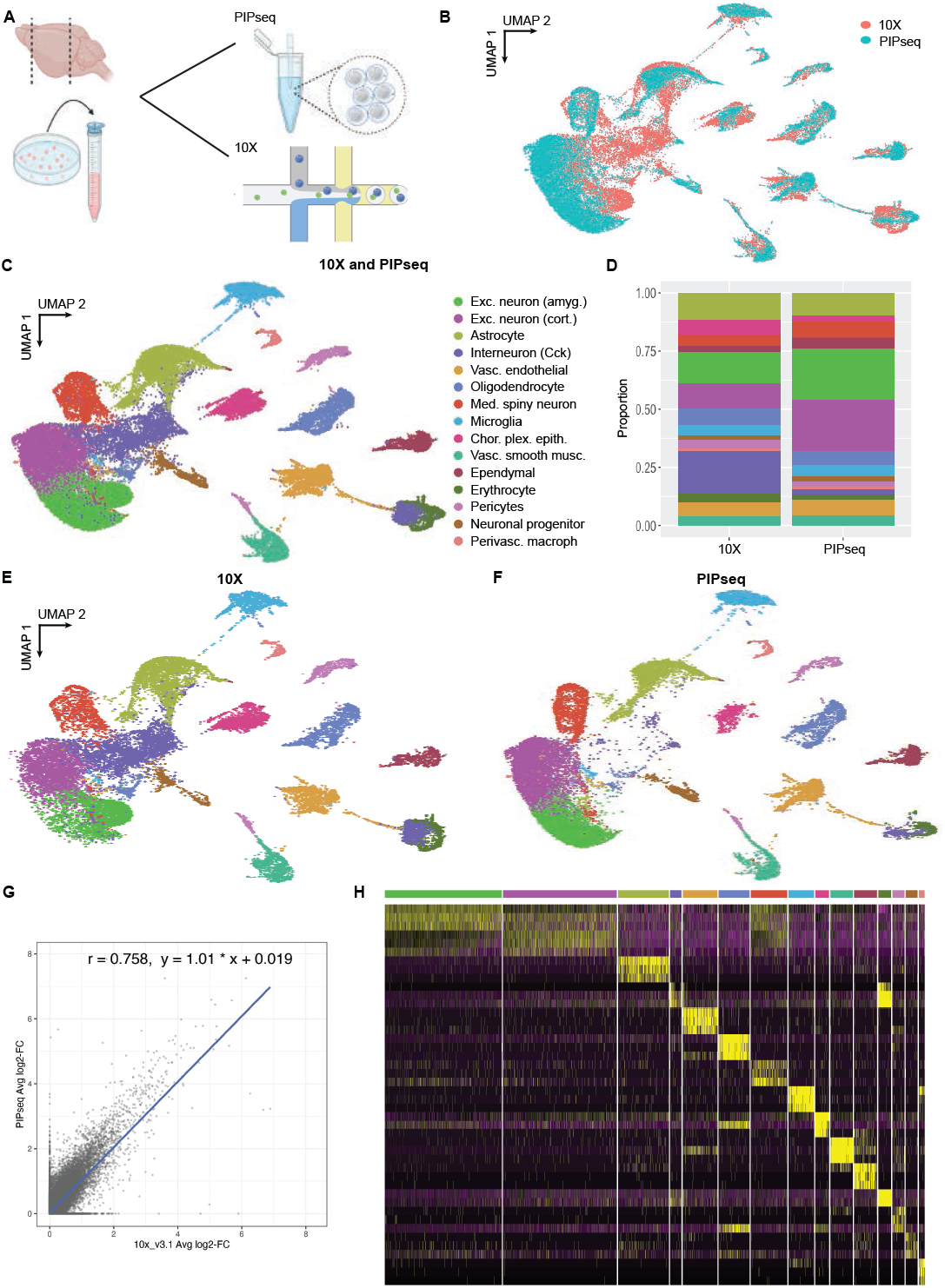
PIPseq analysis of dissociated mouse brain produces comparable data to 10X Genomics. **A)** Experimental setup. A single cell suspension was prepared from P30 mouse brain and was loaded concurrently into pre-templated instant partitions (PIPs) or a 10X microfluidics chip. **B)** Uniform Manifold Approximation and Projection (UMAP) plot of normalized datasets from both single-cell sequencing platforms. Integration was performed using Harmony. **C)** UMAP plot of combined datasets labeled with cell types automatically assigned using scType; see Fig S1C for cell type composition breakdown by sequencing platform. **D)** Proportions of automatically identified cell types in each downsampled dataset; see Fig S1C for cell type composition by sequencing platform and Fig S2 for feature plot data supporting cell type categories. **E**,**F)** UMAP plot of datasets from Fig 1C labeled by sequencing platform origin, 10X (E) or PIPseq (F). **G)** Correlation plot of top differentially expressed genes in 10X vs. PIPseq datasets. **H)** Heatmap of top 5 marker genes for each automatically identified cell type from Fig 1C.

Noting these differences, we next sought to integrate the two datasets and directly compare each platform’s capacity for capture and annotation of different central nervous system (CNS) brain cell types. We used Harmony^20^ to integrate the two datasets (Fig 1B), and used scType^21^ to automatically identify and annotate major CNS cell types using a custom-built reference with curated^12^ cell type markers (Fig 1C; Fig S1C). Calculated cell type marker genes included canonical cell marker genes, including Snca for cortical excitatory neurons, Aldoc for astrocytes, and *Plp1* for oligodendrocytes, among others (feature plots for these genes are found in (Fig S2)). We compared proportions of each cell type identified across the two datasets (Fig 1D-F), and found that all identified cell types were broadly present across both platforms (Fig 1D), considering that cell-type proportion data are difficult to compare across conditions and samples in scRNAseq datasets^4,22^. We did observe a lower proportion of the “R-LM border Cck interneurons” cluster in the PIPseq sample (Fig 1D); however, we noted that marker genes for this cluster and a cluster identified as “Unknown” by scType (cluster 12) included hemoglobin genes such as *Hba-a1*. Plotting *Hba-a1* expression across clusters identified by scType (Fig S3A) confirmed higher expression in the interneuron cluster (cluster 4) relative to background levels, with even higher *Hba-a1* expression noted for the “unknown” cluster (cluster 12). Cluster 12 was subsequently labeled as erythrocytes given its increased *Hba-a1* expression relative to other clusters (Fig S3A). Finally, we plotted the average expression of all hemoglobin genes across all cells from each dataset (Fig S3B), which showed enrichment in 10X data relative to PIPseq for some hemoglobin genes (*Hba-a1, Hbb-bt*) but not others (*Hba-a2, Hbb-bs*). *Hba-a1* expression across each cluster split by single-cell platform (Fig S3C) showed consistently higher *Hba-a1* levels in the 10X dataset across all clusters.

After investigating cell type capture rates from primary mouse brain, we next wanted to compare the relative expression of marker genes between both platforms. Differential expression analysis (Wilcoxon rank-sum) was performed separately for each of the integrated datasets, and the average log_2_-fold change (log_2_fc) was plotted for each detected marker gene. A strong Pearson correlation was observed (r^2^ = 0.7936) between marker enrichment values calculated for each gene across each platform (Fig 1G). Importantly, the top five marker genes detected by 10X for each cell type were robustly enriched in each of the corresponding cell types captured by PIPseq, and their gene expression was used to construct a heatmap that demonstrates cluster-specific expression across the combined (PIPseq + 10X integrated) dataset (Fig 1H).

Thus, PIPseq and 10X have similar performance and capture very similar biological information when using the same mouse primary brain sample. Both techniques capture major brain cell types at comparable rates, with minimal cell-type-specific capture differences across the two scRNAseq platforms. Finally, both approaches can be used to identify marker genes that are enriched in biologically meaningful clusters, a key use for single-cell sequencing technologies across disciplines.

### Large scale PIPseq can efficiently capture subtypes of inflammation-responsive brain cells

A major benefit of PIPseq technology over microfluidic-based scRNAseq platforms is scalability – particularly for the sequencing of large numbers of cells from a single sample. Previous large-scale PIPseq reactions have been reported using cryopreserved human blood cells and banked human reduction mammoplasty specimens^18^. Given these earlier applications in non-nervous tissue, we next applied PIPseq at large scale to map cell identity in the mouse brain using freshly purifi ed cells. CNS tissue is a good test case, as it is a diffi cult tissue to analyze accurately with other scRNAseq techniques – particularly for CNS resident immune and glial cells^15^. Specifi cally, we were interested in using largescale PIPseq to investigate the global changes that occur in CNS cells to systemic infl ammatory insults, as we previously reported low capture rates for astrocytes using this model and the 10X genomics platform^6^.

First, we evaluated whether PIPseq could detect gene expression changes in CNS cells after exposure to a potent infl ammatory agent, the bacterial cell wall endotoxin lipopolysaccharide (LPS). We prepared a single-cell suspension of primary forebrain from two male and two female *Aldh1l1*^*eGFP*^ mice injected peripherally with either LPS (5 mg/kg) or saline, and captured single cells for RNA sequencing using PIPseq (Fig 2A). In each sample, 200,000 cells were added to v3 PIPs following the manufacturer’s instructions. 20,000-50,000 cells per sample were recovered after reverse transcription, library prep, and sequencing for a fi nal dataset of 126,463 cells. Barcoded reads were assigned to cells with PIPseeker, then fi ltered for quality based on visual inspection of outliers (see Methods). Next, the Scanpy software package^23^ was used to scale and normalize gene expression, and data integration/batch correction was performed with Harmony to account for sex differences (Fig 2B). Cells were clustered based on gene expression using standard algorithms (Fig 2C), and then assigned cell types based on expression of canonical marker genes (Fig S5A-D).

**Figure 2.**
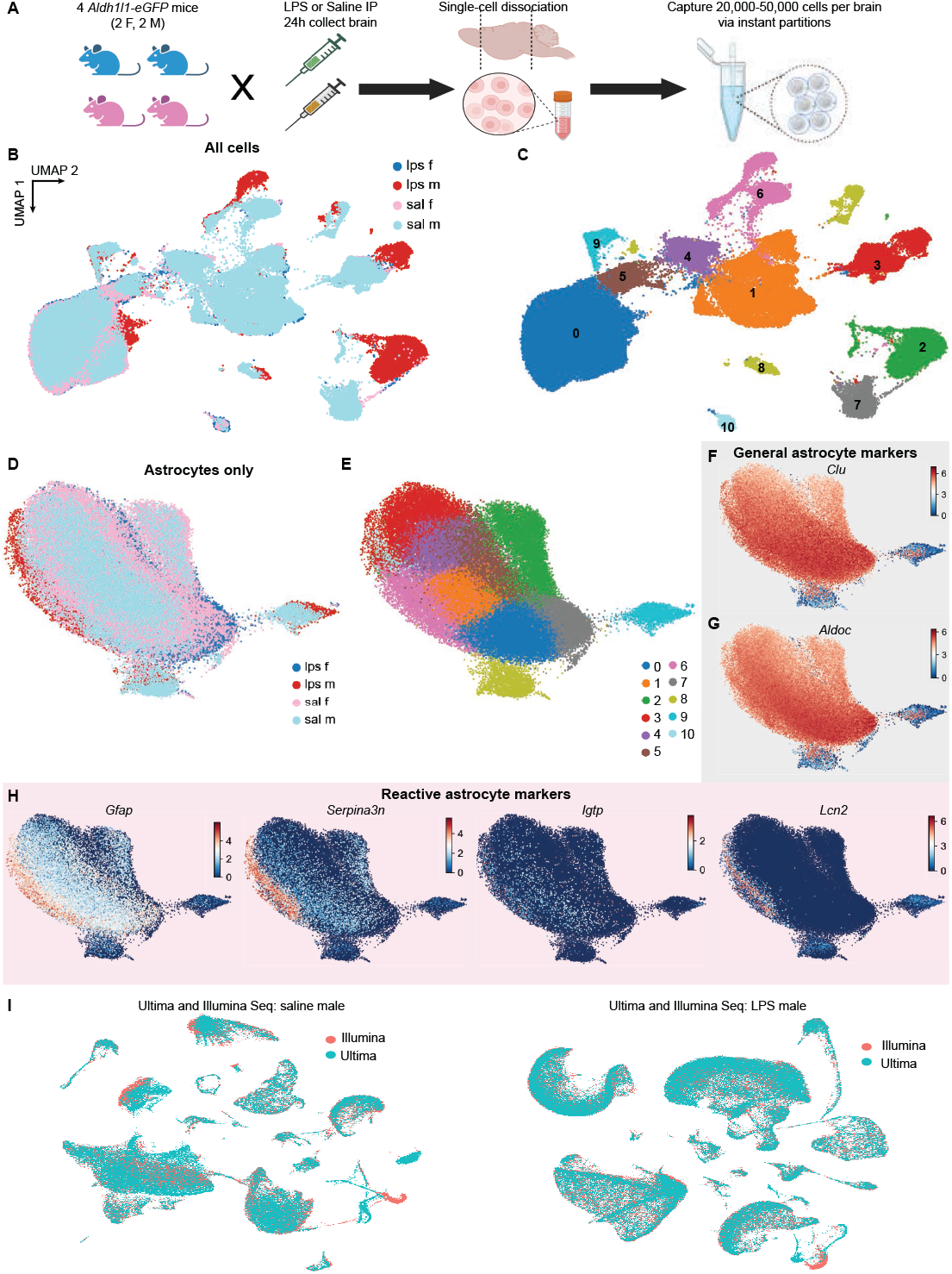
PIPseq analysis at large scales detects differential gene expression in astrocytes in an inflammation mouse model. **A** Experimental setup. Single-cell suspensions were prepared from four mice either treated with lipopolysaccharide (LPS; inflammation) or saline (control) for 24 hours. **B)** Uniform Manifold Approximation and Projection (UMAP) plot of all four samples after clustering; integration was performed using Harmony, quality control information can be found in Fig S4A-C. **C)** UMAP plot of combined datasets labeled with unsupervised cluster labels; see text and Fig S5 for cluster marker gene information. **D)** Astrocytes alone (n = 69,345) were subset from Fig 2C and analyzed as a separate dataset. **E)** Astrocytes were re-clustered in order to find LPS-driven inflamed reactive astrocytes. **F**,**G)** Feature plots for astrocyte marker genes *Clu* (F) or *Aldoc* (G) demonstrate robust expression across entire astrocyte dataset. **H)** Feature plots of reactive astrocyte marker genes show enrichment for these genes in cells from the LPS samples. **I)** Comparison of PIPseq libraries from the saline male sample (left) or LPS male sample (right) sequenced with two different sequencing platforms, Illumina or Ultima. After downsampling to match sequencing depth across platforms and integration using Harmony, there were very minimal differences observed when the same sample was sequenced on the two different platforms; detailed statistics on each sample are in Fig S4D-F.

We reported previously that large numbers of astrocytes are required to identify lowly abundant, spatially restricted, and biologically important reactive astrocyte substates^6,24,25^. Due to cost limitations, this has historically been achieved by pre-enrichment for astrocytes/astrocyte nuclei using FACS/ FANS for the cell type of interest prior to sc/snRNAseq. In contrast, PIPseq has a scalable microfluidic-free technology that allows for capture and sequencing of many more cells from the same sample without the need for FACS/FANS, while still maintaining high numbers of astrocytes. To this end, the astrocytes (clusters 0,5,9 from Fig 2C, 69,345 cells total; see Fig 2F,G) from the LPS dataset were reclustered to look for reactive subpopulations in the LPS experiment (Fig 2D,E). Analysis of cluster marker genes uncovered a clear cluster of 6,332 astrocytes (cluster 6) that highly expresses reactive markers^6^ like *Serpina3n* and *Lcn2* (Fig 2H).

Given the high sequencing costs required to analyze modern, large-scale scRNAseq datasets, particularly with scalable technologies that allow hundreds of thousands of cells to be captured from a single sample, we next examined whether our PIPseq libraries could be sequenced using these alternative sequencing techniques, in line with recent reports^26^. Aliquots of two PIPseq libraries were sequenced using Ultima technologies’ spinning-wafer high-throughput platform, resulting in 3.1 and 2.2 billion reads per sample (Fig S4D). The Ultima reads were also processed using PIPseeker, and merged Seurat objects with cells sequenced using both platforms were used to probe for any sequencing-platform-based differences (Fig 2G; Fig S4E,F). After integrating using Harmony, we saw no appreciable differences in cell clustering due to the sequencing platform (Fig 2G), thus demonstrating that PIPseq can be successfully used with high-throughput sequencing platforms when analyzing large datasets.

### PIPseq can detect differential gene expression across biological conditions

Confident that we could capture large numbers of CNS cells without loss of individual cell types and with appropriate sequencing depth, we next sought to determine if subtler perturbations often used in neuroscience could be detected using PIPseq. To do this, we next tested whether PIPseq has sufficient sensitivity to detect changes in gene expression between transgenic mouse models of different genotypes. The Clusterin/Apolipoprotein J knockout mouse^27^ (*Clu*^*-/-*^) was used as a model system to test PIPseq’s sensitivity for two reasons. First, this gene is highly expressed by astrocytes^19^, and thus is a good test case to examine whether PIPseq could detect the reductions in gene expression expected with a heterozygote or full knockout mouse. Second, recent reports suggested cryptic expression of Clu from an alternate exon in the *Clu*^*-/-*^ mouse^28^, yet no groups have confirmed this observation either using purified astrocytes and bulk RNA-seq, or at the scRNAseq level.

Single-cell forebrain suspensions were prepared from 4 *Clu*^*-/-*^ (knockout/KO), 2 *Clu*^*+/-*^ (heterozygote), and 2 WT C57Bl/6 (ie. *Clu*^*+/+*^) animals at age P32 and 40,000 cells per sample were loaded into PIPs (Fig 3A). After manual data quality control and filtering, Seurat was used to normalize expression data and reduce dimensionality resulting in a dataset of 58,087 cells (WT: 14,687; *Clu*^*+/-*^ Het: 13,871; *Clu*^*-/-*^ KO: 29,529; Fig 3B-D). Harmony was used to correct for sex differences in gene expression before performing differential gene expression (DE) analysis using Seurat (Fig 3E-G). As expected, *Clu* was one of the most highly DE genes across genotypes, detected as a top DE both when comparing WT versus *Clu*^*-/-*^ and WT versus *Clu+/-* across all cells (Fig 3E). Strikingly, in the homozygous *Clu* knockout there was reduced but still extant Clu expression in many cells (Fig 3E), providing evidence across nearly 30,000 cells for recent claims that this mouse line is not actually a true knockout^28^. We next turned to DE analysis in the heterozygote, which appears to be a suitable genetic model for reduced *Clu* expression (Fig 3F). Astrocytes specifically (cluster 1) were tested for other genes that varied with Clu genotype in Clu heterozygotes (Fig 3G; Fig S7); newly discovered DE genes included elements of the ubiquitin-proteasome pathway such as Ubb and related genes. Specific alteration of gene expression of members of the proteasome pathway in the brains of Clu knockout mice have not been reported previously, although Clu plays a major role in proteostasis in many other systems^29,30^. Interestingly, there were reciprocal increases in the expression of the Ubb-ps pseudogene in clusters 5 and 12 in *Clu*^*+/-*^ animals (Fig S8A,B), in agreement with the reported inverse relationship of expression levels between some pseudogenes and their reciprocal gene^31^. Overall, PIPseq was able to detect changes in gene expression in cells from a knockout mouse model, including both alterations in *Ubb* and further evidence for persistent (yet severely decreased) expression of *Clu* in the putative knockout line.

**Figure 3.**
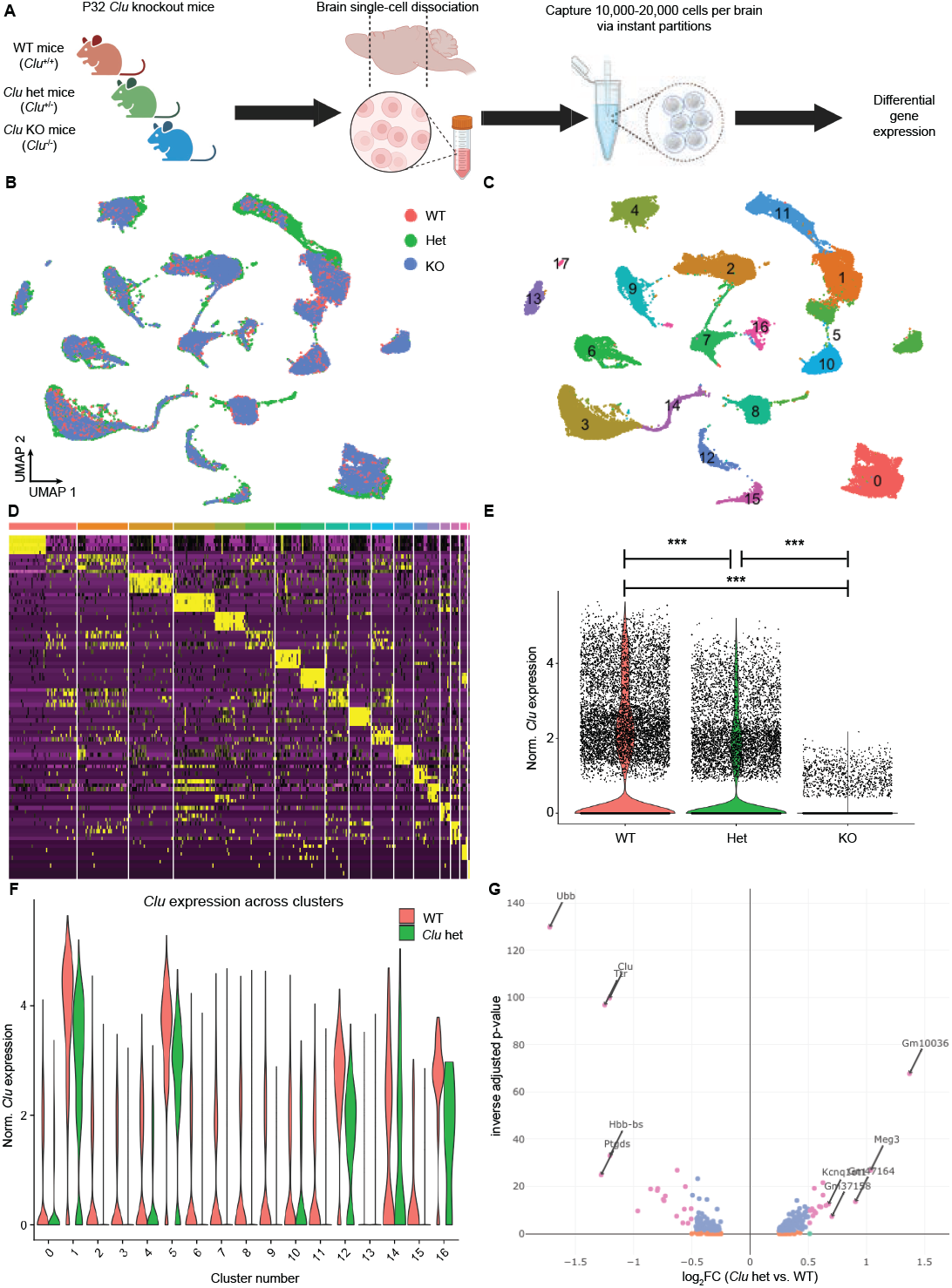
PIPseq analysis detects differential gene expression in an genetic knockout mouse model. **A)** Experimental setup. Single cell suspensions were prepared from P32 Clusterin (*Clu*) knockout mouse brains and 20,000 cells per sample were loaded into pretemplated instant partitions (PIPs). **B)** Uniform Manifold Approximation and Projection (UMAP) plot of all cells (n = 58,087) after clustering and integration for batch correction using Harmony. **C)** UMAP plot of the full dataset labeled with cluster IDs; see Fig S7A for cluster ID labels at low resolution, and see Fig S7C-E for marker gene expression plots supporting cell type assignments. **D)** Heatmap of top 5 marker genes for each cluster from Fig 3C. **E)** Violin plot of normalized Clu expression in all cells from each genotype shows significantly lower *Clu* expression in heterozygote and KO vs. wild-type astrocytes (p < 0.001; Wilcoxon ranked-sum test. n= 14,687 cells, 2 animals [wild-type]; n= 13,871 cells, 2 animals [heterozygote]; n= 29,529 cells, 4 animals [knockout]). However, low-level but clear *Clu* expression is noted in the putative KO, confirming recent reports of cryptic *Clu* expression in this supposed knockout animal (see text). **F)** Violin plot of *Clu* expression in WT vs. heterozygote animals, plotted by cluster IDs from Fig 4C. *Clu* is highly expressed in astrocyte clusters 1,5 and other cells (cluster 12, 14, 16). **G)** Volcano plot of genes differentially expressed in Clu heterozygotes vs. WT cells from only astrocytes (cluster 2) in Fig 4C. Clu is significantly downregulated in heterozygotes vs. WT, in addition to Ubb, a gene linked to Clu function. See Fig S8 for other volcano plots for other clusters.

Next, PIPseq was used to detect virally-induced transgenes, a common genetic tool for cell subtype targeting in neuroscience and biological research more broadly. To test this, we dissected hippocampi from four *Ptenflx/flx* X Ai14D^*TdTomato*^ transgenic mice, two of which were injected with control lentiviruses (FUGW) and two that received lentivirus with *Cre* recombinase (FUG-T2A-Cre; Fig 4A). Upon receiving viral *Cre*, transgenic mice from this cross should 1) lose *Pten* expression and 2) begin to express *TdTomato* under a CAG promoter. Nuclei extraction was performed with a simple single tube commercial kit (see Methods), and 40,000 nuclei per sample were loaded into PIPs. cDNA was produced and sequenced following manufacturer’s instructions, then barcoded reads were mapped to a custom transcriptome containing the tdTomato gene (see Methods; quality control data in Fig S9A). 101,491 nuclei were recovered in total across the four animals, and then clustering and dimensionality reduction was performed as previously demonstrated (Fig 4B-D; Fig S9). Cell types were called based on marker gene expression (Fig S10) and compared with published dentate gyrus scRNAseq datasets^32^. Expression of the *TdTomato* transgene was detected in 129 nuclei from the two mice (63,690 nuclei total) that received the FUG-T2A-Cre virus (Fig 4E). There was also tdTomato expression in 3 nuclei from the two FUGW injected animals (37,933 nuclei total), likely as a result of background, Cre-independent expression of the transgene^33^. Thus, PIPseq can detect expression of tdTomato in transgenic mice after induction via viral *Cre*. This capture rate represents less than 0.2% of all nuclei captured, a number of cells likely too lowly abundant to be captured using other sc/snRNAseq methods without prior enrichment using FACS/FANS.

**Figure 4.**
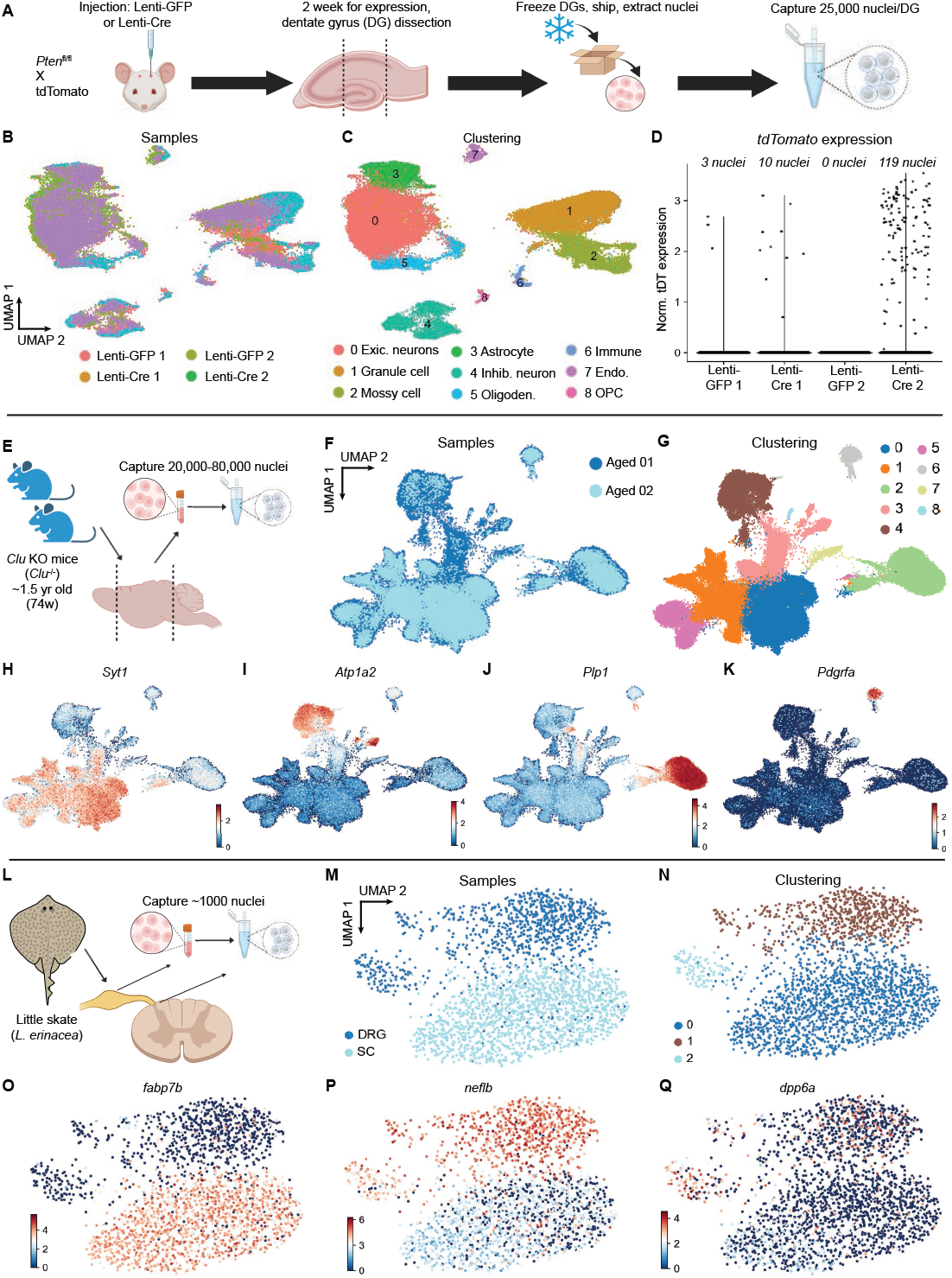
PIPseq can be used to sequence nuclei from diverse tissues and experimental conditions. **A)** Experimental setup for panels A-D (dentate gyrus dataset). *Pten*^*flx/flx*^ x TdTomato^*flx/flx*^ mice were injected with either Lenti-GFP (control) or Lenti-Cre viruses; the *Cre* lentivirus should delete *Pten* expression and induce *TdTomato* expression. After waiting 14 days for viral expression, dentate gyri were dissected from 2 control and 2 Cre mice, then nuclei were extracted and processed with PIPseq. **B)** Uniform Manifold Approximation and Projection (UMAP) plot of all nuclei (101,491) after clustering and integration for batch correction using Harmony. **C)** UMAP plot of the full dataset labeled with cluster IDs; marker gene expression data can be found in Fig S10C-E. **D)** Violin plot of *TdTomato* expression by sample demonstrates clear enrichment of the transcript (log_2_fc: 3.702, p < 0.001) in the two mice that received Lenti-Cre (Lenti-Cre 1 and Lenti-Cre 2). *TdTomato* expression in the control samples was similar to *Xist* expression in males (e.g., low to nonexistent and thus likely background; Fig S9C). **E)** Experimental setup for panels E-K (aged mouse nuclei dataset). Nuclei were extracted from whole forebrain of two aged (74 week old) *Clu* knockout mice (see Fig 3), and processed with PIPseq. **F)** UMAP plot of all nuclei (104,729) after clustering and integration. **G)** UMAP plot of all nuclei labeled by cluster; marker gene feature plots are in following panels. **H)** Feature plot of neuronal marker gene (*Syt1*). **I)** Feature plot of astrocyte marker gene (*Atp1a2*). **J)** Feature plot of oligodendrocyte marker gene (*Plp1*). **K)** Feature plot of oligodendrocyte precursor marker gene (*Pdgfra*). **L)** Experimental setup for panels L-Q (skate spinal cord nuclei dataset). Nuclei were extracted from the spinal cord or dorsal root ganglia of multiple little skate (*Leucoraja erinacea*) embryos at equal developmental stages. 5,000 nuclei per tissue were processed with PIPseq to pilot the nuclei extraction from this tissue and test bioinformatic resources in this non-traditional model species. **M)** UMAP plot of all nuclei (DRG: 951, SC: 1,485) after clustering and integration. DRG = dorsal root ganglion, SC = spinal cord. **N)** UMAP plot of all nuclei labeled by cluster; marker gene feature plots are in the following panels. **O)** Feature plot of *fabp7b* expression (spinal cord marker gene). **P)** Feature plot of *neflb* expression (previously validated DRG marker gene). **Q)** Feature plot of *fabp7b* expression (cluster 2 marker gene).

We next examined Pten expression in the 132 *TdTomato* positive nuclei, which was decreased relative to GFP-injected controls (Fig S9B). Nuclei that received the lentiviral *Cre* and thus were positive for tdTomato should have zero *Pten* expression due to excision of the *Pten* gene; however, Pten expression was >0 in 21/132 total TdTomato+ nuclei analyzed (Fig S9B). This observation could represent 1) a subset of cells that are misclassified as TdTomato positive, 2) persistent expression of *Pten* from a non-floxed exon in the knockout animal, similar to reports from other transgenic strains^28^, or 3) contamination from ambient RNA, leading to background like that seen in the minute level of *Xist* expression seen in male nuclei (Fig S9C).

Overall, these two experiments using global genetic ablation (*Clu*) and virally mediated deletion (*Pten*) demonstrate that PIPseq can accurately detect and quantify changes in gene expression across biological conditions, in both common and rare cell types.

### PIPseq allows capture and profiling of aged brain nuclei

PIPseq was next tested on nuclei isolated from aged mouse brain samples, a problem of particular interest given that changes in cell-type-specific gene expression are reported in many CNS cells throughout aging^17,34,35^. Extraction of nuclei from cells is an important use case for single-cell analysis, as nuclei are often the preferred input for pathology samples (often obtained as fresh-frozen blocks from tissue banks) or for aged tissue that resists simple enzymatic dissociation^17^. Nuclei were isolated from fresh-frozen aged (∼P518) *Clu*^*-/-*^ (*Clu* KO) mouse brain through a combination of physical and chemical isolation methods, notably without the use of enzymes. With this simple nuclei prep, 160,000 dissociated nuclei were loaded into PIPs as outlined above, followed by standard library prep and sequencing (Fig 4E,F). Aged nuclei data were analyzed using the same normalization and dimensionality reduction strategy as was applied to whole cells, resulting in 104,729 nuclei after outlier exclusion (Fig S11A). Nuclei were then clustered and classified into cell types based on marker gene expression (Fig G-K). All CNS cell types were identified in the aged nuclei dataset (as in the sc/snRNAseq data from younger mouse brains – see Fig 2C, Fig 3C). Taken together, these data demonstrate that PIPseq robustly and efficiently captures primary nuclei from the aged mouse brain in a large scale (100,000s of nuclei) single-batch reaction, and produces data of sufficient quality to clearly identify and cluster brain cell types.

### PIPseq allows capture and profiling of nuclei from spinal cord in the little skate

To extend the analysis of PIPseq efficacy beyond mouse brain, nuclei were extracted from dorsal root ganglion (DRG) and spinal cord (SC) from the little skate (*Leucoraja erinacea*). To verify that the genomic resources for this nonmodel species would be sufficient for further snRNAseq experiments and analysis, we took advantage of the low-input requirement for PIPseq to run a pilot experiment on low numbers of nuclei. 5,000 nuclei from each tissue were loaded into PIPs (Fig 4L), and after outlier exclusion a dataset of 1,485 nuclei from SC and 951 nuclei from DRG remained (Fig 4M). Clustering at low resolution (0.2) revealed 3 distinct clusters (Fig 4N), which could be defined by expression of cluster specific genes (*fabp7b* for SC, *neflb* for DRG; Fig 4O-Q). The observation of *neflb* expression in the DRG and SC samples is partially validated by past work showing specific expression of this gene in zebrafish SC by in situ hybridization^36^, in addition to DRG-specific enrichment of mouse *Nefl* in a mouse DRG scRNAseq dataset^37^. Although *neflb* expression in the skate data is clear, the DRG dataset displayed reduced genes per nucleus and counts per nucleus compared to the SC sample (Fig S11B-C), even though the DRG sample was sequenced more deeply (DRG: 43,338 reads/nucleus, SC: 17,704 reads/nucleus). Thus, this lower depth may represent a biological difference, or alternatively a bioinformatic difference in gene annotation level between the two tissues, given the non-model reference genome. To conclude, even at low sequencing depth in a pilot experiment setting, PIPseq recovered major cell types from nervous system tissues other than brain in a non-traditional model species.

## DISCUSSION

The sequencing of individual cells and nuclei from the CNS has unearthed myriad novel cell types, sub-states, and putative cell-cell interactions during development, aging, and in disease. Many of these discoveries have been enabled solely by the advent of commercial single cell technologies

– often at a cost that is prohibitive to many end users, or not achievable if core facilities are not present at a particular institution. Here we employed microfluidics-free, scalable, low-cost PIPseq to investigate gene expression in single cells and nuclei in the brain across a variety of experimental approaches and questions.

First, PIPseq and 10x datasets produced from the same sample were directly compared, with minimal differences found between the platform. This finding of minimal difference is in line with the original PIPseq paper, which performed a similar comparison between immortalized cells^18^. Concurrently, every major cell population from a difficult-to-dissociate primary tissue (brain) was detectable with both platforms, suggesting that both approaches may be used to produce datasets with comparable cell capture. We note the many technical complexities that make it difficult to measure and compare cell capture rates across individual scRNAseq experiments^38^; these complexities similarly make it difficult to precisely compare cell capture across different capture technologies (e.g., microfluidic vs. non-microfluidic).

We produced both large scale (up to 100,000) and small scale (∼1,000) single cell and nucleus datasets using PIPseq, with a combination of both Illumina and Ultima sequencing. Very little differences in downstream data analysis were noted when comparing the same PIPseq library sequenced across both platforms. Given the potential for PIPseq to eventually achieve single-batch million-cell capacity for scRNAseq capture^18^, sequencing costs will quickly become the limiting variable when planning experiments; thus, our demonstration of compatibility with newer sequencing techniques suggests that million-cell experiments may ultimately be attainable at reasonable cost for research groups of many sizes. On the other end of the spectrum, our small scale (∼1,000 cell) spinal cord experiment shows the efficacy and ease of using PIPseq to run small pilot experiments. This capacity allows easy, low-cost piloting in experiments where sample prep (single-cell/nuclei dissociation) may take multiple rounds of optimization, and is similarly useful in nontraditional model organisms, where a researcher might not know a priori if extant bioinformatic resources suffice to run a single-cell experiment. Thus, PIPseq produces comparable datasets to microfluidics-based single-cell technologies like 10X, and PIPseq libraries are able to be sequenced by multiple independent sequencing platforms.

In conclusion, this study provides a first test-case for the use of low cost, microfluidics-free, and scalable PIPseq for the investigation of nervous system tissue across age, brain regions, experimental application, and species. In contrast to the high barrier to entry, rigid sample size, and high cost of other commercial systems, PIPseq can produce comparable and fully integratable sequencing data that should enable rapid piloting (at low input) and validation (at high input) of many neuroscience and biological research questions.

## METHODS

### Experimental model details

#### Mice (Mus musculus)

This study used young (P32) and aged (P350) C57B/6J mice, *Aldh1l1*^*eGFP*^ mice ((Tg(Aldh1l1-EGFP)OFC789Gsat/Mmucd, RRID:MMRRC_011015-UCD), *Clu*^*-/-*^ mice (Clu<tm1Jakh>,RRID:IMSR_JAX:005642), and *Pten*^*flx/flx*^ (RRID:IMSR_JAX:004597) crossed to the Ai14^*TdTomato*^ Cre reporter mouse (RRID:IMSR_JAX:007914). All mice were housed on a 12-h light/dark cycle with food and water ad libitum. All animal procedures were in accordance with the guidelines provided by the National Institute of Health as well as NYU Langone School of Medicine’s Administrative Panel on Laboratory Animal Care. All animals were housed at 22–25 °C and 50–60% humidity. All procedures at Dartmouth were approved by Dartmouth College’s Institutional Animal Care and Use Committee (IACUC) and Association for Assessment and Accreditation of Laboratory Animal Care Review Board. Animals of either sexes were utilized for these experiments.

#### Little Skate (Leucoraja erinacea)

Embryos at the desired developmental stages (stages 32+/33) of the little skate, *Leucoraja erinacea*, were obtained from the Marine Resources Department of Woods Hole Marine Biology Laboratory (MBL). The egg cases were kept in seawater prepared with Instant Ocean (Aquarium Systems) at a temperature of 16 °C using a standard aquarium filtration system. Prior to dissection, the embryos were anesthetized with 1X Tricaine (MS-222, Sigma-Aldrich) diluted with seawater.

#### Modelling systemic inflammation

To model an inflammatory response, the bacterial cell wall endotoxin lipopolysaccharide (LPS, Sigma, L2880) was injected intraperitoneally (5 mg kg^-1^) and animals euthanized for cell collection 24 h later as previously described^7^. Control animals received an injection of equal volume of endotoxin-free 0.9% saline (Enzo, ALX-505-009-LD15).

#### Viral tracing

7-day old (P7) *Pten*^*flx/flx*^ x Ai14 animals were stereotaxically injected with 2 μL amount of virus in the dentate gyrus following published protocols^39^. Brains were dissected in sterile PBS 14 days after injection, and successful injection was confirmed under a fluorescence microscope. Dentate gyri were dissected and frozen, then shipped overnight on dry ice for nuclei extraction, PIP capture, processing and sequencing at NYU.

### Method details

#### Cell and nuclei isolation

*Mouse:* For mouse cell dissociation, whole postnatal day (P) mouse forebrains were dissociated according to a modified papain-based protocol^19^ (Worthington), then filtered through a 40 μM filter before counting and dilution for scRNAseq. Viability was estimated at 90-95% based on trypan blue exclusion while counting cells.

For mouse aged brain nuclei dissociation, aged (∼P350) mice were sacrificed with CO_2_ and their brains were immediately dissected and flash frozen in liquid nitrogen. Nuclei were then extracted using a modified commercial kit (Sigma Nuclei EZ Prep, #NUC101-1KT). Mouse forebrain was dissected, and each hemisphere was separately flash-frozen in liquid nitrogen. Frozen hemispheres were thawed, diced with razor blade, then placed in a dounce homogenizer with 1 mL EZ for 20 strokes loose, 20 strokes tight. The dounce was washed with another 1 mL of EZ solution, brought to 5 mL EZ on ice and left to sit for 5 min. Tubes were spun at 600 rcf for 5 min at 4 °C then EZ buffer was removed, nuclei were washed with 5 mL PBS plus 0.1% BSA and RNAse inhibitor then spun again. Nuclei were resuspended in 50-500 μL PBS to count.

*Little Skate:* Skate spinal cords and dorsal root ganglia (DRGs) were extracted from embryos in seawater at stage 32+/33 and collected in 500 μL of filtered sEBSS (∼45 mL EBSS with 1 g Urea, 0.7 g NaCl, and 0.2 g TMAO). The collected samples were centrifuged at 300 × g for 4 min at 4°C to remove the supernatant, rapidly frozen using dry ice, and stored at -80 °C.

Spinal cords from three embryos and DRGs from six embryos were used to collect 5,000 nuclei. Frozen samples were transferred to a chilled dounce for mechanical homogenization in 2 mL of cold EZ Lysis Buffer (similar to mouse nuclei extraction). The samples were then incubated on ice for 5 min, centrifuged at 500 × g for 5 min at 4 °C, the supernatant was discarded, and the samples were resuspended in cold EZ lysis buffer. These steps were repeated, and the samples were resuspended in cold Nuclei suspension buffer (5 mL PBS, 50 μL NEB BSA (10 mg/mL), and 25 μL RNase inhibitor (40 U/μL)). The samples were centrifuged, resuspended in ∼1 mL of cold NSB once more, filtered through a 40-μm cell strainer, and centrifuged again. Finally, the samples were resuspended in 20 μL of cold Cell suspension buffer. 5,000 nuclei were then loaded into PIPs, cDNA was synthesized and libraries prepared following manufacturer’s instructions. The two samples were sequenced on an Illumina NextSeq2000 flowcell, with a goal of 25,000 reads/nucleus (actual: DRG: 43,338 reads/nucleus, SC: 17,704 reads/nucleus).

#### Single-cell and single-nucleus capture via PIPs

Single cells were partitioned into PIPs following manufacturer’s instructions. Briefly, 40,000-100,000 live cells or nuclei were added to a tube containing template beads thawed on ice, pipetted gently to mix, and then immediately vortexed for 175 seconds and processed further according to manufacturer’s instructions. For some experiments, stable PIP emulsions containing captured mRNA after cell lysis were shipped on frozen ice packs to Fluent Biosciences (Watertown, MA, USA) for cDNA generation and amplifications, cDNA purification, library preparation and sequencing, performed on a NextSeq 2000 (Illumina).

For 10X genomics comparison experiments, ∼34,000 cells per sample were loaded onto a 10X chip G (overloaded to try to reach parity with PIPseq numbers) using v3.1 3’ chemistry. Reverse transcription and library preparation were performed according to manufacturer’s instructions, and sequenced at the NYU Genome Technology Center (RRID: SCR_017929) using a NovaSeq 6000 sequencer (Illumina). Resulting sequenced libraries were downsampled to the read depth discussed in the text using the seqtk package (v1.2-r94).

#### Ultima Sequencing

Illumina-compatible libraries were shipped to Ultima Genomics for library conversion and sequencing. Samples were sequenced to a depth of ∼2-3 billion reads, as described elsewhere^26^, producing single-end reads with a mean read length of ∼250bp. Prior to processing with PIPseeker, FASTQs were preprocessed to transform the reads into the R1/R2 format used as input by PIPseeker. Read preprocessing included trimming of the polyT, and transposase adapter sequences, and barcode sequences were matched to the barcode whitelist, allowing for a single substitution or indel. R1 was created by concatenating the corrected barcodes with the expected linkers. R2 was created as the reverse complement of the sequence following the polyT, discarding 5 bases following the polyT. Final libraries from each sequencing platform were downsampled to equal read depths as discussed in text.

### Quantification and statistical analysis

#### Data processing and quality control

To analyze the 10X Genomics datasets, .fastq files from sequencing were demultiplexed, then used as input for Cell Ranger (v6.0.1) for assignment of reads to single cells. PIPseq datasets were analyzed using PIPseeker (v1.0.0, then v2.1.4 for the skate data), a comprehensive analysis solution that performs barcode matching, quality control, mapping with the STAR aligner^40^ deduplication, transcript counting, cell calling, clustering, differential expression and cell type annotation. scType (v1.0) was used for automatic cell calling following recommended instructions and code from that package, with a reference annotated dataset from mouse brain^12^. For *Pten* experiments, a custom STAR mouse reference genome was built by adding the tdTomato gene to mouse genome version mm10-2020-A.

Datasets for Figs 1,3,4 were analyzed in Seurat (v.4.3.0). For the data in Fig 2, visible outliers were filtered in Scanpy (v.1.8.1) according to the following parameters: genes_by_ counts < 4,000, n_counts < 20,000, pct_counts_mt < 35. This led to 126,463 cells passing quality control out of 130,963 cells originally output by PIPseeker. Harmony^20^ v0.1.0 was used to integrate across conditions where noted, to account for batch effects.

#### Skate bioinformatics

The chromosome-level genome assembly and annotated transcriptome of the little skate^41^ was accessed from NCBI (RefSeq: GCF_028641065.1). Chromosome names in the genome were edited to match the transcriptome chromosome names, then the edited .fna and .gtf files were used to make an indexed genome using STAR^42^. This indexed genome was used to process raw .fastq files via PIPSeeker (v2.1.4). Nuclei were excluded from further analysis based on the following cutoffs: n_gene_by_counts <2500, total_ counts <5000 (see Fig S11A for plots of post-filter statistics). DE analysis was performed with a Wilcoxon ranked-sum test in Scanpy, as per above.

### Statistics

Marker genes for each cluster and related differential expression (DE) analyses were calculated using the Wilcoxon rank-sum algorithm implemented in Scanpy or Seurat, and clustering was performed using the Louvain algorithm. Volcano plots for DE were made using the Plotly package (v4.10.1). All statistical tests used are highlighted in the legend of each figure. No data were excluded from the analyses, except when performing quality control filtering as discussed above. The experiments were not randomized. The investigators were not blinded to allocation during experiments and outcome assessment.

## Acknowledgements

The computational requirements for this work were supported in part by the NYU Langone High Performance Computing (HPC) Core’s resources and personnel. We also acknowledge the shared resources of the NYU Langone Genome Technology center that is partially supported by a Cancer Center Support Grant P30CA016087 at the Laura and Isaac Perlmutter Cancer Center. Funding for this work was provided by the Cure Alzheimer’s Fund, MD Anderson Neurodegeneration Consortium, Paul Slavick, the NIH/NEI R01EY033353, and The Alzheimer’s Association (SAL); NINDS T32 (NYU/Dasen) 5T32NS086750 (PWF); NIMH R01MH097949 (BWL); PIPseq development at Fluent BioSciences was supported in part by NIH/NIGMS 1R43GM137648 and 1R44GM145185. Some graphics were made in BioRender.

---

**Supplementary Figure 1.**
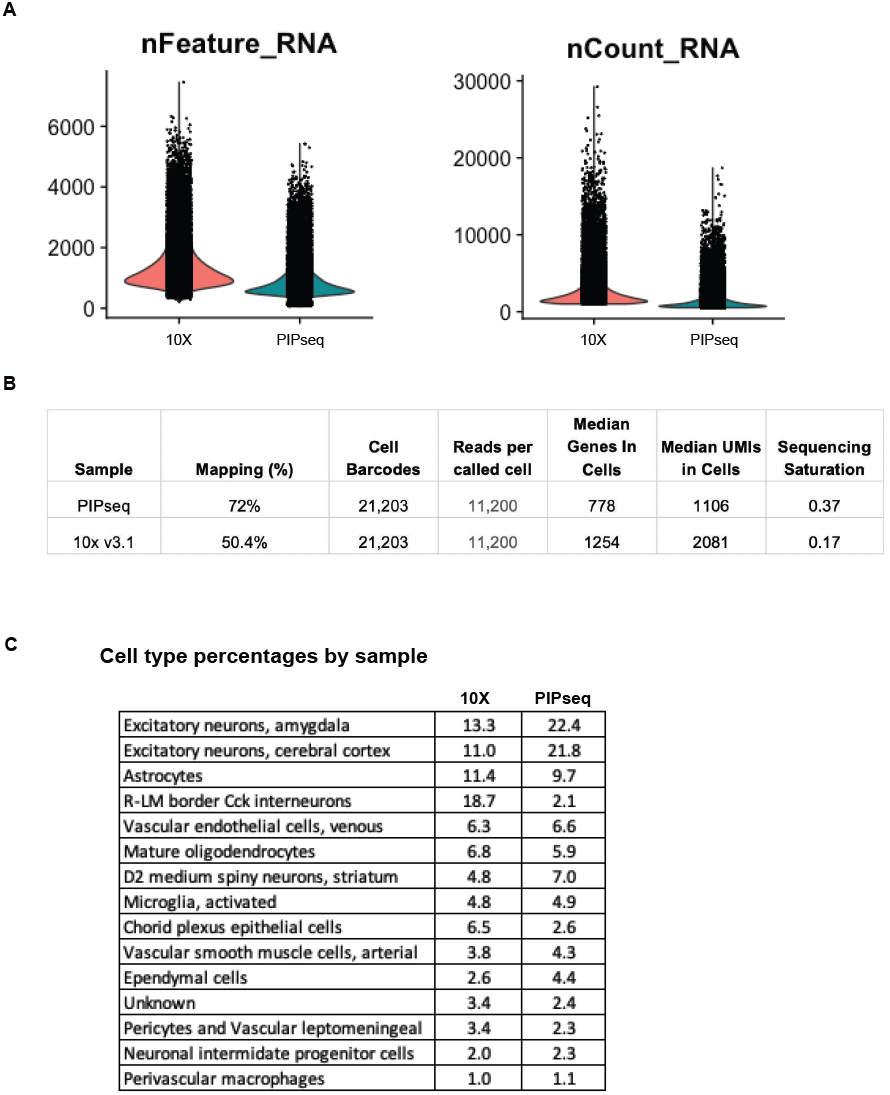
Quality control and sequencing information for Fig 1. **A)** Plot of genes call per cell in combined, downsampled dataset (Fig 1B-H). **B)** Mapping and other quality control statistics for downsampled dataset. **C)** Table of percentages of each automatically identified cell type found in each sample of the combined dataset. Percentages are listed out of total cells from each respective single-cell sequencing platform (24,562 cells for 10X and 21,203 cells for PIPseq).

**Supplementary Figure 2.**
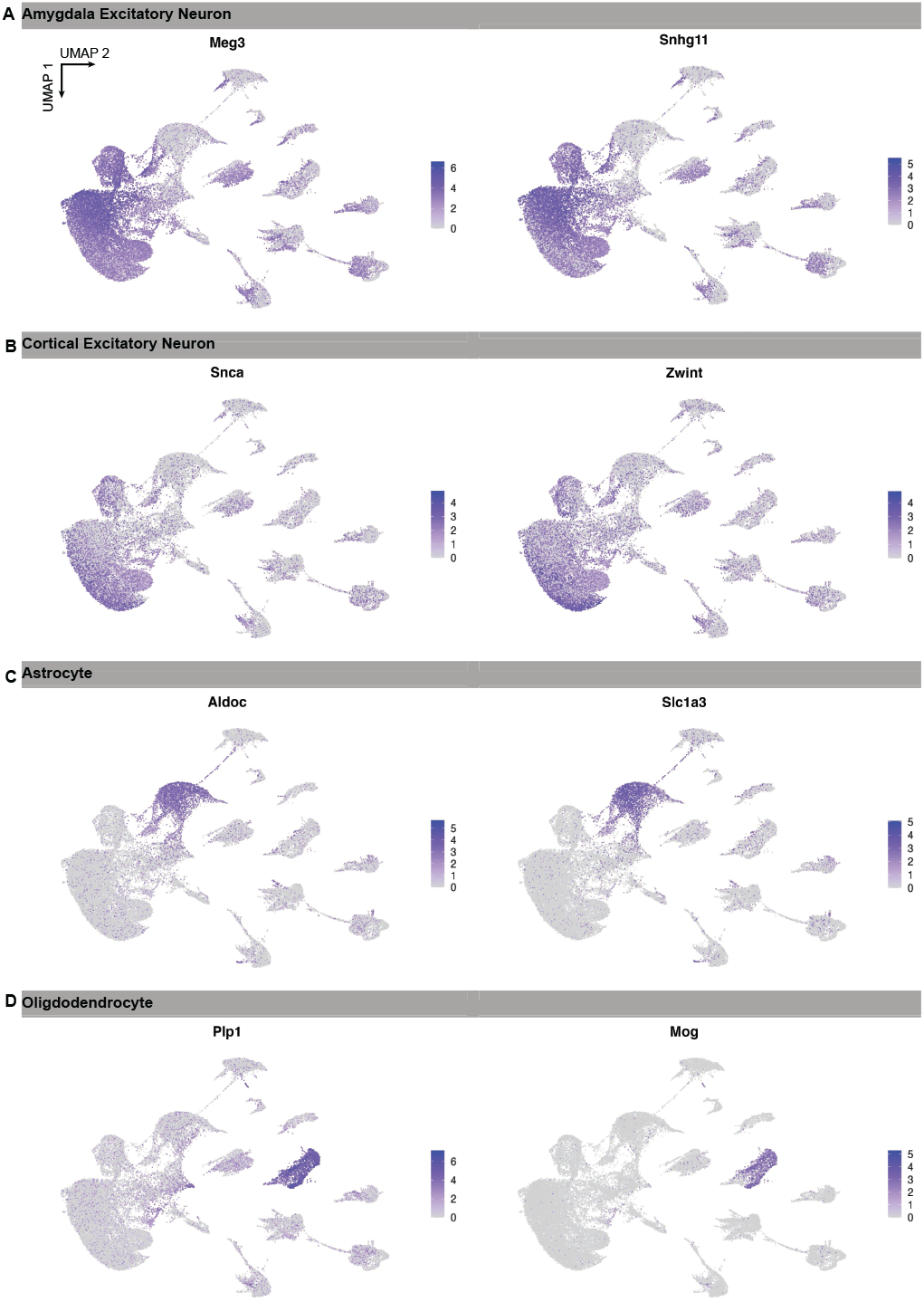
Gene expression supporting automated cell type calling in Fig 1. **A)** Feature plots of marker genes for amygdala excitatory neurons (*Meg3* and *Snhg11*). **B)** Feature plots of marker genes for cortical excitatory neurons (*Snca* and *Zwint*). **C)** Feature plots of marker genes for astrocytes (*Aldoc* and *Slc1a3*). **D)** Feature plots of marker genes for oligodendrocytes (*Plp1* and *Mog*).

**Supplementary Figure 3.**
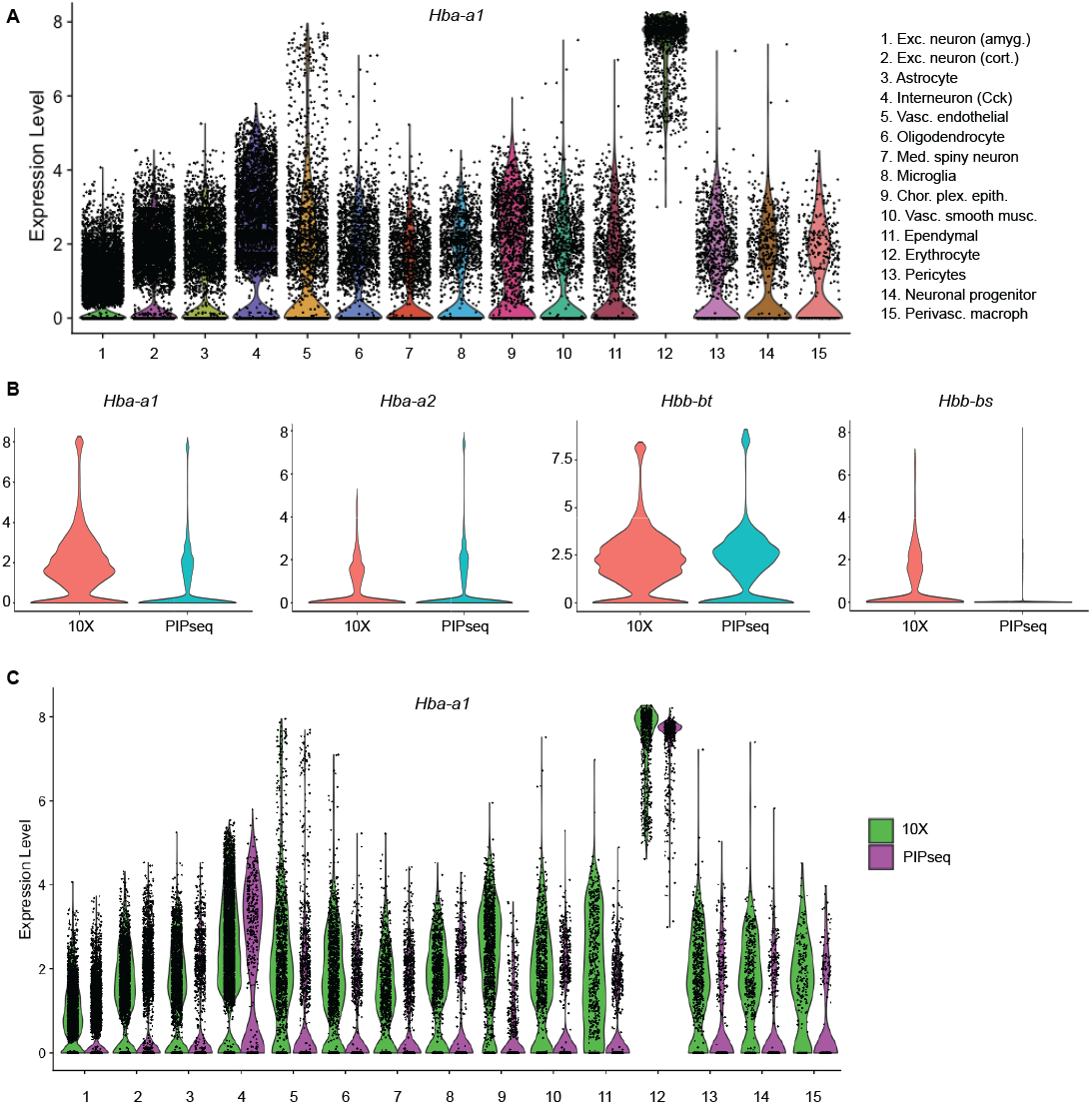
Hemoglobin gene expression analysis across platforms. **A)** *Hba-a1* expression across all automatically-identified cell types. Clear enrichment is noticed in cluster 12. **B)** Violin plots of gene expression for four hemoglobin genes split by scRNAseq platform. Expression is consistent across platforms, except for *Hbb-bs* which seems enriched in the 10X dataset. **C)** Similar expression pattern of *Hba-a1* across all cell types, split by scRNAseq platform.

**Supplementary Figure 4.**
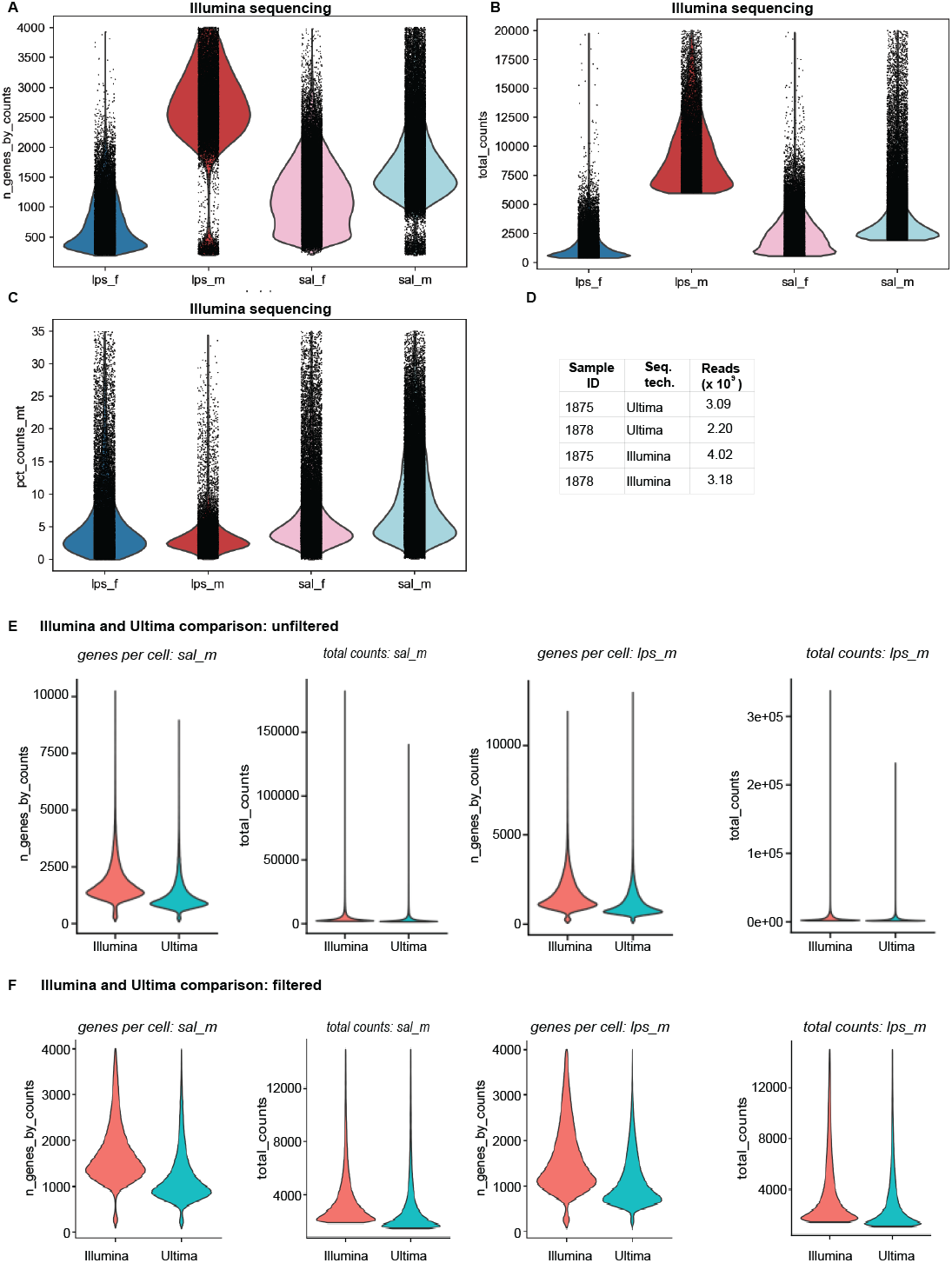
Quality control and sequencing information for Fig 2. **A-C)** Quality control information for four samples from Fig 2. **D)** Table of sequencing reads generated per sample from each sequencing platform (Illumina or Ultima). **E-F)** Quality control graphs for both samples combined for each sequencing platform for all cells (unfiltered, E) or the subset analyzed in Fig 2I (filtered, F).

**Supplementary Figure 5.**
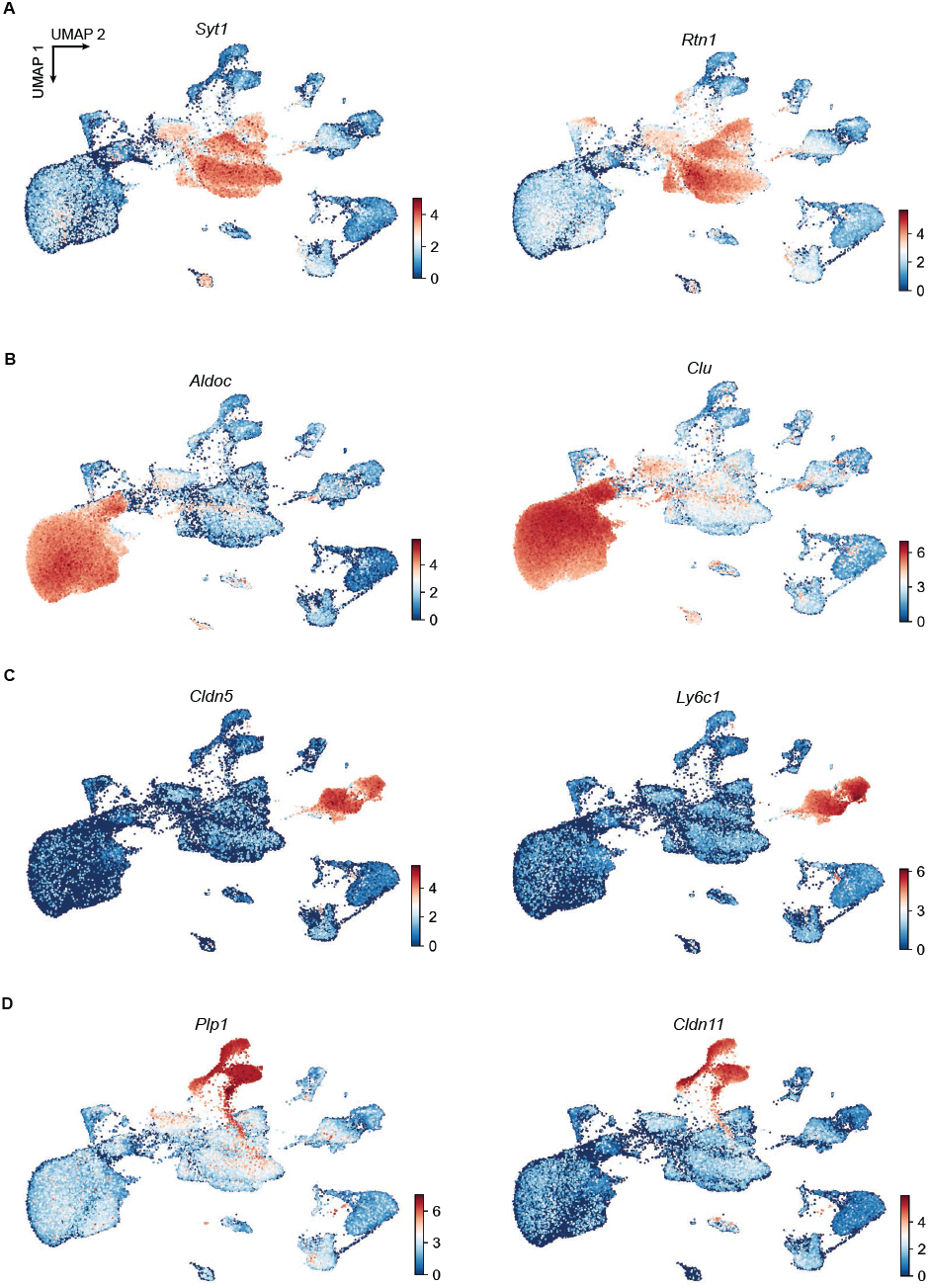
Gene expression supporting cell type calling in Fig 2. **A)** Feature plots of marker genes for cluster 1 from Fig 2C (neurons; *Syt1* and *Rtn1*). **B)** Feature plots of marker genes for cluster 0 from Fig 2C (astrocytes; *Aldoc* and *Clu*). **C)** Feature plots of marker genes for cluster 3 from Fig 2C (endothelial; *Cldn5* and *Ly6c1*). **D)** Feature plots of marker genes for cluster 6 from Fig 2C (oligodendrocytes; *Plp1* and *Cldn11*).

**Supplementary Figure 6.**
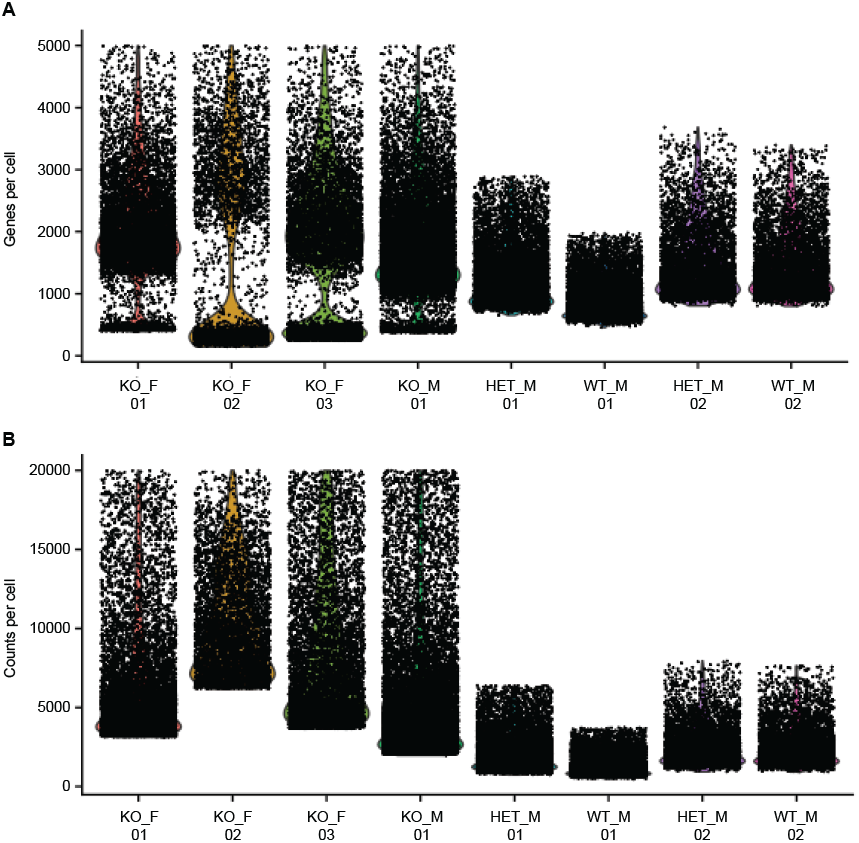
Quality control and sequencing information for Fig 3. **A)** enes per cell for each sample from Fig S6. KO= knockout (*Clu*^*-/-*^), Het= heterozygote (*Clu*^*+/-*^), WT= wild type (*Clu*^*+/+*^) B) Counts per cell for each sample from Fig S6.

**Supplementary Figure 7.**
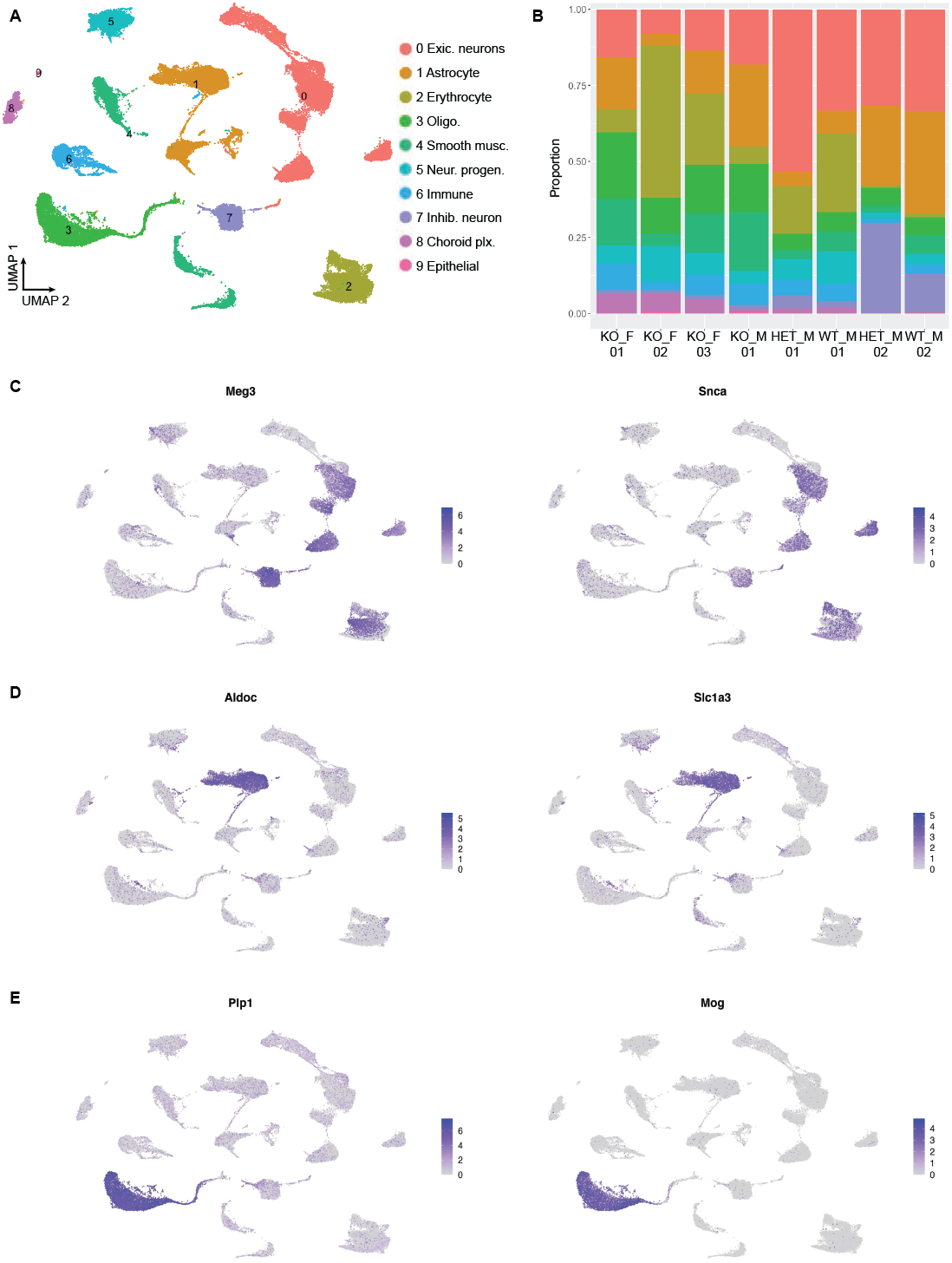
Gene expression supporting cell type calling in Fig 3. **A)** Dataset from Fig 3C resclustered at low resolution. Clusters are labeled based on expression of cell type marker genes (Fig S7C-E). **B)** Plot of cluster proportions recovered from each individual animal from Fig 3. **C)** Feature plots of marker genes for cluster 0 from Fig S7A (excitatory neurons; *Meg3* and *Snca*). **D)** Feature plots of marker genes for cluster 1 from Fig 2C (astrocyte; *Aldoc* and *Slc1a3*). **E)** Feature plots of marker genes for cluster 3 from Fig 2C (oligodendrocytes; *Plp1* and *Mog*).

**Supplementary Figure 8.**
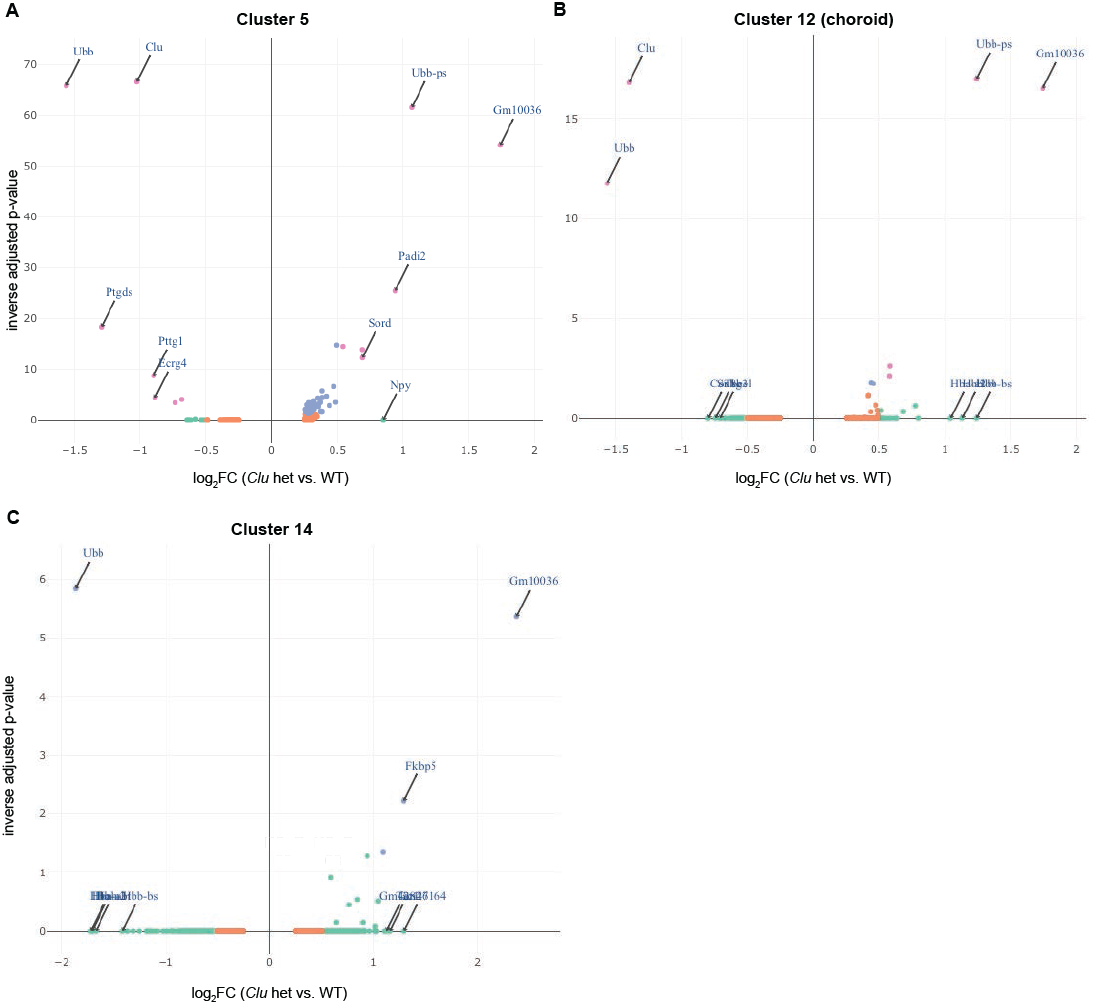
Extended differential gene expression (DE) analysis for Fig 3. **A)** Volcano plots of genes DE in *Clu* heterozygotes vs. WT controls for cluster 5 from Figure 3 (astrocytes #2). DE analysis was performed using a Wilcoxon-ranked sum test implemented in Seurat, see Methods for full details. DE genes include *Ubb* and *Ubb-ps* as discussed in the text, plus *Clu* itself as expected. **B)** Volcano plots of genes DE in *Clu* heterozygotes vs. WT controls for cluster 12 from Fig 3 (choroid plexus cells). **C)** Volcano plots of genes DE in *Clu* heterozygotes vs. WT controls for cluster 12 from Fig 3 (ependymal cells).

**Supplementary Figure 9.**
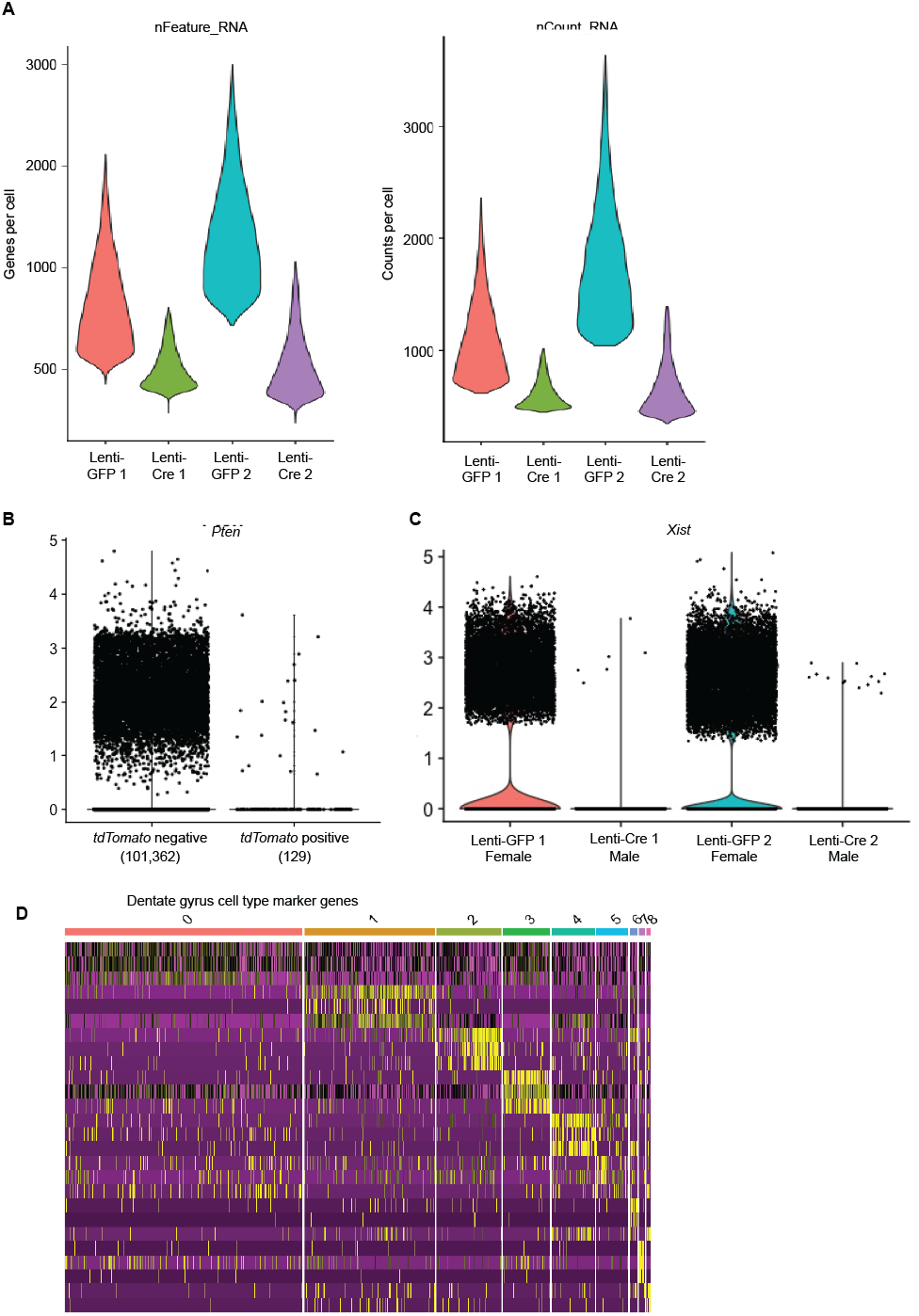
Quality control and sequencing information for dentate gyrus data (Fig 4A-D). **A)** Genes per cell and counts per cell for each of the four dentate gyrus (DG) samples from Fig 4A-D. **B)** Violin plot for Pten expression in cells that did not express / *TdTomato* (left) and in *TdTomato* positive cells (right). **C)** Violin plot for *Xist* shows comparable background expression to *Pten* expression in *TdTomato* positive cells (previous panel). **D)** Heatmap of top 5 marker genes for each DG cluster from Fig 4C.

**Supplementary Figure 10.**
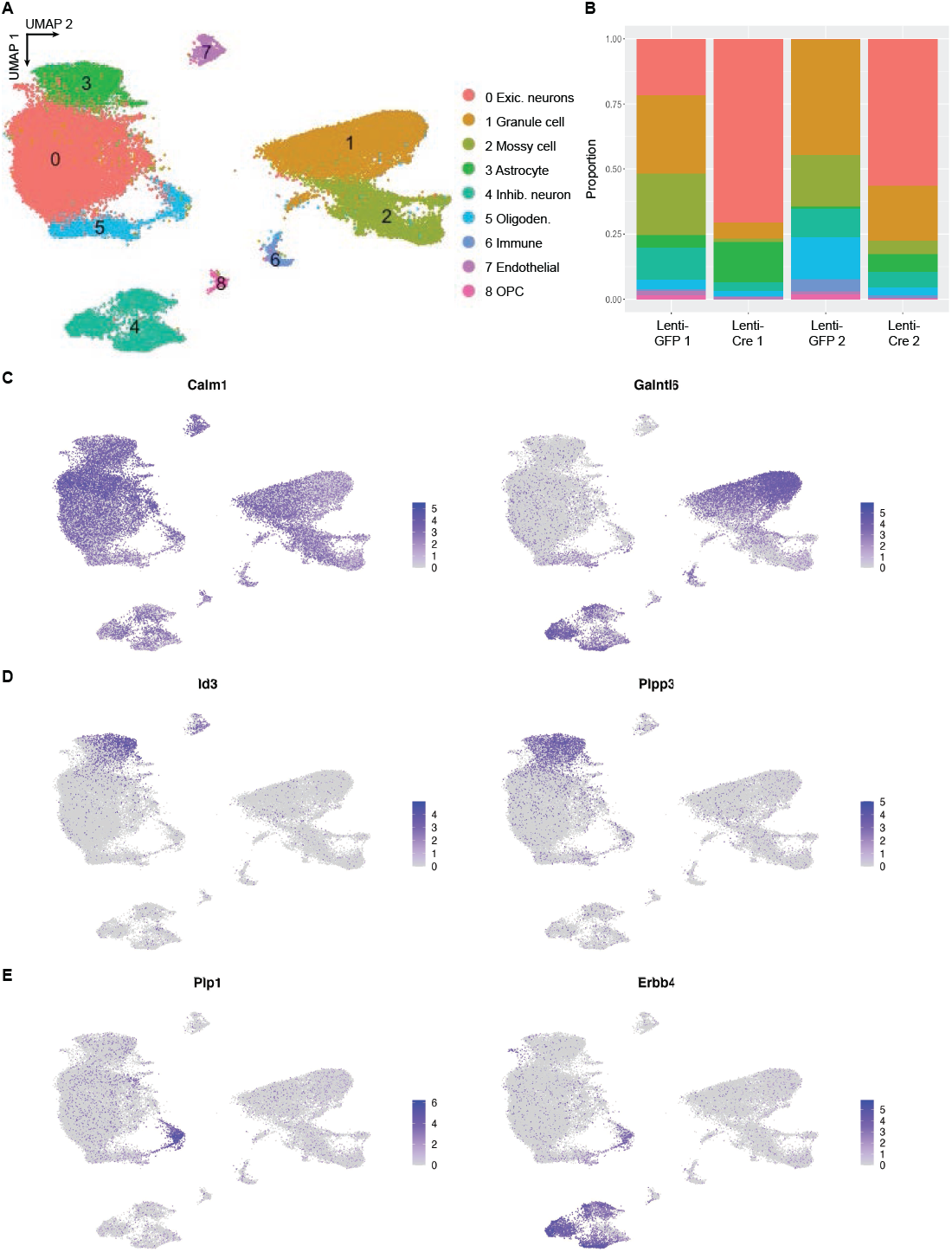
Gene expression supporting cell type calling for dentate gyrus data (Fig 4A-D). **A)** Data from Fig 4C replotted for ease of marker gene comparison below. **B)** Cell type proportion graphs from each animal. **C)** Feature plots of marker genes for cluster 0,1,2 from Fig 4C (neurons; *Calm1* and *Galntl6*). These genes were previously reported as dentate gyrus markers in^32^. **D)** Feature plots of marker genes for cluster 3 from Fig 4C (astrocytes; *Id3* and *Plpp3*). E) Feature plots of marker genes for cluster 5 from Fig 4C (oligodendrocytes; *Plp1*).

**Supplementary Figure 11.**
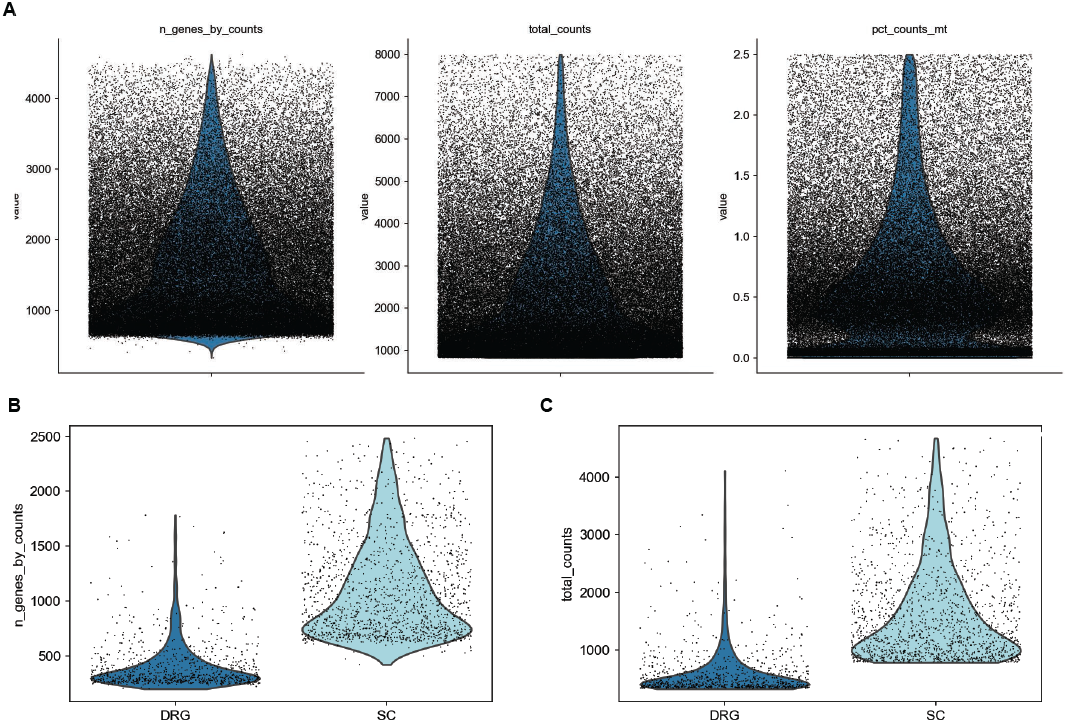
Quality control and sequencing information for aged mouse and skate data (Fig 4E-Q). **A)** QC data (genes/ nucleus, counts/nucleus, mitochondrial count/nucleus) for both samples combined from the aged mouse dataset, Fig 4E-K. **B)** QC data for little skate (*Leucoraja erinacea*) nuclei pilot dataset, Fig 4L-Q, split by sample (DRG: dorsal root ganglia, SC: spinal cord).

